# ARG1 could be expressed by human, but not represent for repair

**DOI:** 10.1101/2020.07.02.183624

**Authors:** Chenyu Chu, Chen Hu, Shengan Rung, Yufei Wang, Yili Qu, Yi Man

## Abstract

Arginase 1 (ARG1), typically expressed under stress or stimulation rather than in the healthy condition in humans. However, more and more studies claim ARG1 could be expressed by human and represents healing process. Including THP-1 monocyte shift to M2 macrophages rendered a higher expression of ARG1 than that with bone-graft has an ideal effect on anti-inflammation, resulting in tissue healing. With the higher expression of ARG1 seemingly induced by the constructed material and represented by the anti-inflammatory situation. For these reasons, this study focuses on whether traditional markers of macrophages could be express by human or not during clinical healing process. Through harvesting bone tissue samples during bone healing process and confirm the expression of markers at transcriptome and protein levels. The results showed the expression of ARG1 relatively low by bulk-RNA sequencing, quantitative polymerase chain reaction (qPCR), immunofluorescence (IF) and flow cytometry or could not be detected during process of healing of healthy donor. Taken together, results indicate that the ARG1 is not chosen for a regenerative marker in human bone tissue.

## 1 Introduction

Certain structures and components have the ability to influence cell-cell interaction and phenotypes of macrophages ^1^. However, there exists a controversial issue in this article regarding an enzyme, arginase 1 (ARG1), as it is typically only expressed under stress or stimulation, such as in patients with tuberculosis ^2^, burn victims ^3^, and in blood cells from those suffering from multiple sclerosis ^4^, rather than in the healthy condition in humans. Previous study showed the THP-1 monocyte shift to M2 macrophages rendered a higher expression of ARG1 than that with extra-fibrillarly mineralized collagen (NEMC), which suggests that intra- fibrillarly mineralized collagen (HIMC) has an ideal effect on anti-inflammation, resulting in tissue healing. For the reasons that higher expression of ARG1 seemingly induced by the constructed material and represented for the anti-inflammatory situation whether ARG1 expression as traditional markers of macrophages could be express by human has become a key determinant of this conclusion. Moreover, it aims to come up with the final determination of macrophage phenotype by combining the expression level of ARG1 with traditional related cytokines, including IL-1β, IL-4, IL-10, nos2, etc. It is very interesting to regard the reason for choosing this marker, as ARG1 is only expressed under the effect of disease and trauma on macrophages, as mentioned above. Additionally, the authors’ description of ARG1 as an “anti-inflammatory” “cytokine” that was the marker of the “M2 macrophage” to “promote tissue healing” focused mainly on the effect of ARG1 without emphasis on species differences; previous studies showing ARG1 as a marker for anti-inflammation and healing in macrophages have been restricted to murine models, and have not studied human models ^5^. Moreover, being an anti-inflammatory indicator, ARG1 will be produced when stimulated by anti-inflammatory interleukins, such as IL-4 and IL-13. In other words, ARG1 would be expressed only in murine rather than in human systems under inflammatory conditions. ARG1 can merely be regarded as an anti-inflammatory indicator in cases of murine inflammation ^6^. In addition, excessive activation of ARG1 could be either protective or destructive; the healing effect does not suit the overall situation, which raised our interest in the exact expression of ARG1 in humans, as understood from *in vivo and in vitro* studies ^7^. After processing healthy human peripheral blood mononuclear cells (PBMC), from the 10× genomics library datasets, through single cell RNA sequencing, and antibody staining for flow cytometry (**Supplemental Figure. 1 and Figure. 2**). We in advance confirmed that ARG1 is not expressed, nevertheless, the relationship between ARG1, macrophage, and its phenotypes in local tissues under different status still needs further exploration. To have a more comprehensive understanding of ARG1, its expression in human tissues and THP-1 monocyte-derived macrophages was analyzed from the perspective of both *in vivo* and *in vitro* studies through bulk RNA-seq, qPCR, flow cytometry analysis, and immunofluorescent staining. The harvest procedure for normal bone, inflammatory, and healed tissues is shown in **Figure 1A**, whereas the *in vitro* cell culturing process can be seen in **Figure 2A** (for details, refer to the experimental section).

**Figure 1.**
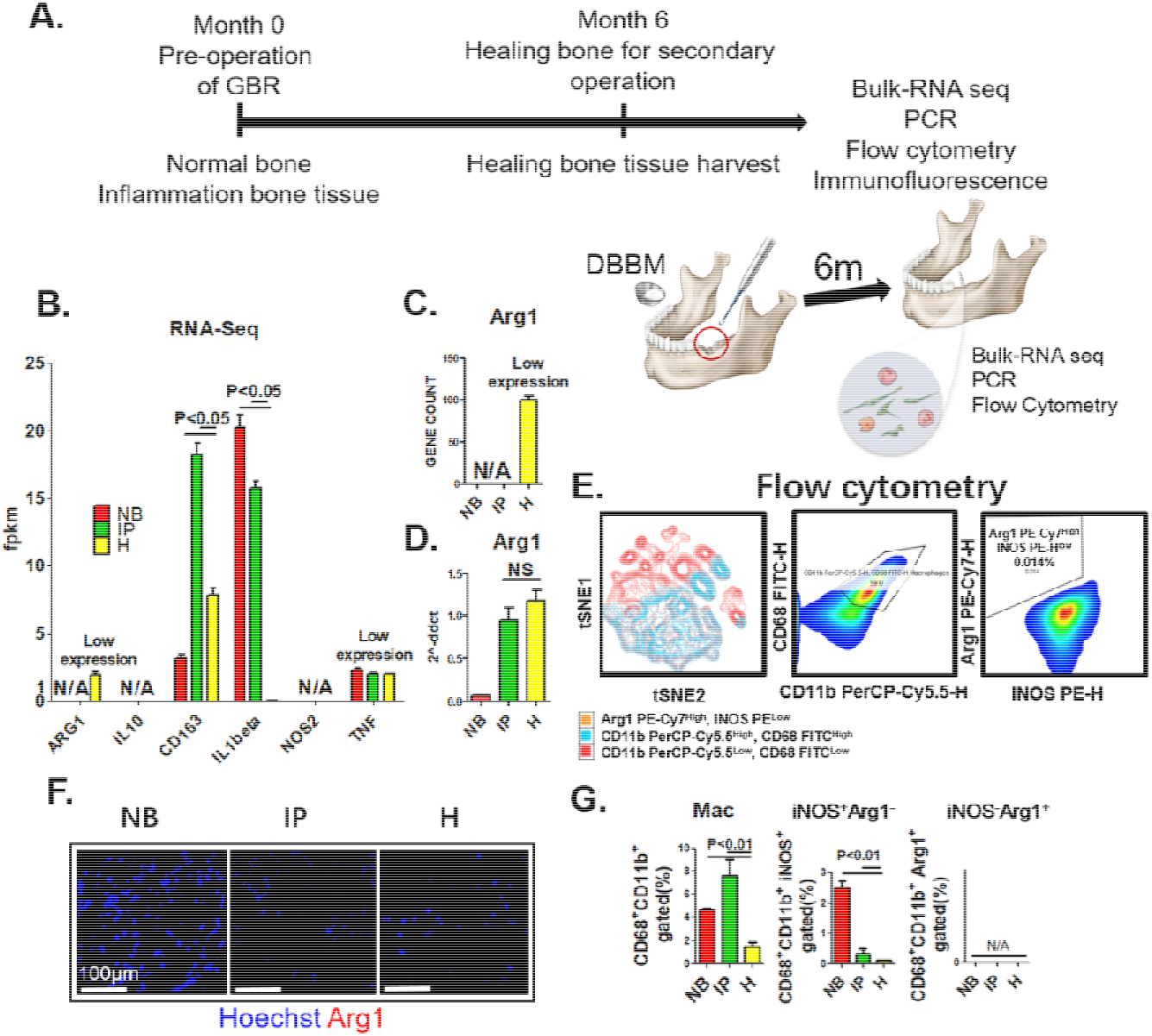
Detection of ARG1 in human normal alveolar bone tissue, inflammatory periodontal tissue, and bone tissue at the healing site. (A) The schematic diagram shows the time point after the pre-operation of GBR when tissues were harvested and the assays performed. (B and C) Analysis of RNA-seq data: both fpkm (B) and gene count (C) confirmed that ARG1 was not detected in the normal and inflammatory states, and its expression was low at the healing site, even when compared with other genes related to macrophages. (D) qPCR assay of ARG1 showed no significant differences between tissues from inflammatory and healing sites. Tissues were collected and digested to obtain single-cell suspensions for flow cytometry analysis. CD68and CD11b were used as markers for myeloid macrophages. iNOS and ARG1 were used as markers for M1 and M2 macrophages. (E) ARG1 expression was not detected in either group, and CD11b^+^CD68^+^ARG1^+^iNOS^−^ macrophages were not observed using gated and t-distributed stochastic neighbor embedding (t-SNE) dimension reduction to visualize high-dimensional data. (F) Immunofluorescent staining did not reveal ARG1 in any group. Hoechst staining appears as blue in color while ARG1 appears as red. (G) Detection of surface markers by flow cytometry analysis. High expression of CD68 (CD68-FITC, Clone Y1/82A) and low expression of Arg1 were evident in the three groups. The population size of CD11b^+^CD68^+^ARG1^−^iNOS^+^ macrophages was very small. Abbreviations: NB, normal alveolar bone tissue; IP, inflammatory periodontal tissue; H, healing (tissue harvest after guided bone regeneration surgery), and N/A, undetected.

**Figure 2.**
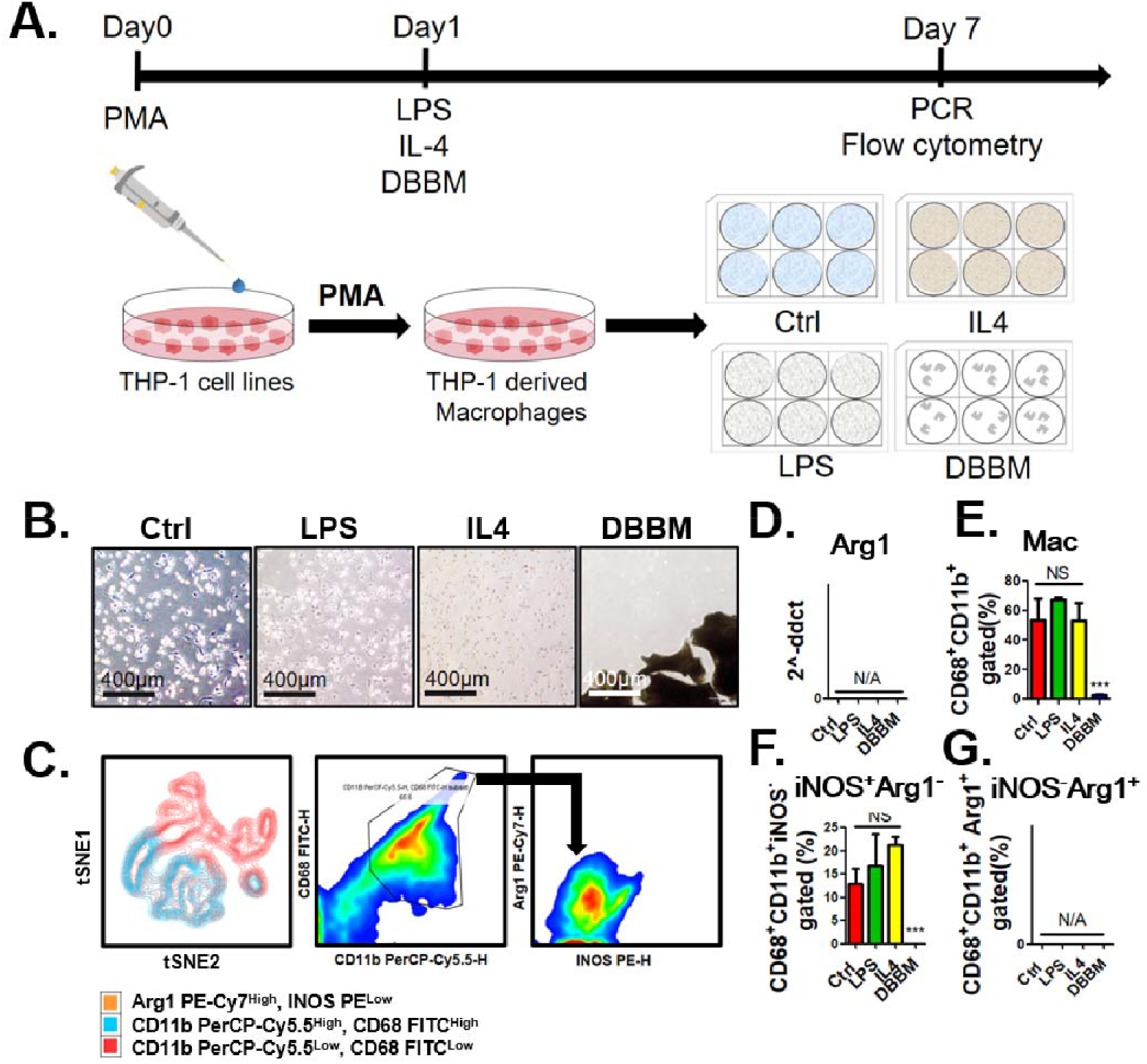
Detection of the ARG1 surface marker after culturing THP-1 monocyte derived macrophages with different stimuli *in vitro*. (A) Schematic diagram of the cell culture process and the time points for stimulation of macrophage differentiation, seeding macrophages in the different conditions, and performing the assays. (B) Light microscopy images of macrophages cultured under different conditions on Day 7. (C) Overlay of tSNE dimensionality reduction diagrams revealed the absence of AGR1^+^ macrophages in the flow cytometry analysis. (D) qPCR assay of ARG1 using Gapdh as the reference gene did not detect ARG1 in the four groups. (E-G) In contrast to the large population of CD11b^+^CD68^+^ macrophages (E) and CD68^+^CD11b^+^iNOS^+^ARG^−^ (F), CD68^+^CD11b^+^iNOS^−^ARG1^+^ macrophages were not detected (G). Abbreviations: Ctrl, RPMI supplemented with 10% FBS; LPS, RPMI supplemented with 10% FBS and 10 ng/ml LPS; IL4, RPMI supplemented with 10% FBS, 10 ng/ml IL4; DBBM, RPMI supplemented with 10% FBS and 0.03 g/well DBBM; NS, no significance; N/A, undetected. ^***^ denotes P < 0.001.

## 2 Combined Results and Discussion

### 2.1 Bulk RNA-seq, quantitative real-time polymerase chain reaction (qPCR), immunofluorescent staining, flow cytometry results of human normal, inflammatory, and healing alveolar bone tissue indicate the insignificance of ARG1, not to mention its relationship with M2 macrophage

A gene count under 10 and fpkm under 1 were considered as not expressed; a gene count lower than 100 was considered as low expression. Based on the above premise, in the normal alveolar bone tissue (NB, control), the gene count showed a relatively low expression of the selected gene, while ARG1 seemed undetected either with the perspective of gene count or fragments per kilobase of transcript per million mapped reads (fpkm). The similar results were observed in the bulk RNA-seq results (**Figure. 1B** and **Figure.1C**) in the other groups, inflammatory periodontal tissue (IP, inflammation) and healing bone tissue (H, healed tissue) harvested at 6 months post-surgery. Moreover, it was considerably lower than the expression level of other macrophage-related genes (**Figure. 1B**). The qPCR results showed the same trend, with no expression in normal bone tissue and no significant differences (P > 0.05) between H and IP tissues (**Figure. 1D**). Detection of ARG1 protein was verified simultaneously. The immunofluorescent staining revealed that ARG1 expression could not be detected among the three groups (**Figure. 1F**). In addition to directly exploring the expression of ARG1 in human tissues, the relationship between ARG1 and macrophage phenotypes is another important objective. From flow cytometry analysis, CD11b^+^CD68^+^macrophages and CD11b^+^CD68^+^ARG1^−^iNOS^+^ M1 macrophages were observed with significant (P < 0.01) differences among the three groups, whereas the nonexistence of CD11b^+^CD68^+^ARG1^+^iNOS^−^ M2 macrophages denied the close relationship between the ARG1 and M2 phenotypes (**Figure. 1E** and **Figure. 1G**). That is to say, both in the inflammatory and in the healthy bone tissue, ARG1 gene expression stays in a relatively static state and is too uncharacteristic to serve as an anti-inflammatory marker in human bone tissue. It is interesting that all our results suggest that the gene and protein expression of ARG1 indicates neither inflammation nor repair status. When focusing on ARG1 expression of specific iNOS^−^ M2 macrophage, negative results in normal, inflammatory, and repaired bone tissue further suggest that ARG1 expression has nothing to do with M2 phenotype.

### 2.2 Arg1 cannot explained the phenotypic classification after conventional stimulation of THP-1 monocyte derived macrophages

Considering that the behavior of cells varied *in vivo* and *in vitro*, we further verified the expression of ARG1 through *in vitro* experiments. The same stimulation method as that described in the discussed article was used to obtain the macrophage from the THP-1 monocyte. Cells were cultured under different conditions; control group (Ctrl), RPMI with 10% FBS; anti-inflammatory group, RPMI with 10% FBS and 10 ng/mL IL-4; inflammatory group, RPMI with 10% FBS and 10 ng/mL LPS, RPMI with 10% FBS; bone graft group, 0.03 g/well deproteinized bovine bone mineral (DBBM). After culturing THP-1 monocyte derived macrophage for 7 days under various conditions (**Figure. 2A-B**), qPCR was implemented, and none of the groups showed a CT value of ARG1 lower than 32 (**Figure. 2D**). Meanwhile, markers including CD11b, CD68, iNOS (NOS2), and ARG1 were stained for flow cytometry analysis. The results exhibited no significant differences in the population size of the CD11b^+^CD68^+^ macrophage among the Ctrl, IL4, and LPS groups; however, it was significantly smaller in the DBBM group (**Figure. 2E**). The number of iNOS^+^ M1 macrophages was also not statistically different among the Ctrl, IL4, and LPS groups, and was much less for the DBBM group (**Figure. 2F**). The CD11b^+^CD68^+^ARG1^+^iNOS^−^ macrophages were not detected in any of the groups (**Figure. 2C and Figure. 2G**). The results showed that when iNOS^+^Arg1^−^ was adopted as M1 macrophage and iNOS^−^Arg1^+^ stood for as M2 macrophage, the former would in accord with the conventional definition of M1 macrophage, whereas the latter failed to explain the inexistence of M2 macrophage after IL-4 stimulation.

In short, both *in vivo* and *in vitro* data confirmed the insignificant expression ARG1 as the marker for human. Moreover, currently, none of the published articles confirm that ARG1 in humans could serve as an ideal marker for repair. It is inappropriate to use ARG1 as a tissue healing biomarker for the M2 phenotype of induced macrophages without experimental evidence. In addition, our results are in support of not using ARG1 to identify M2 macrophages in gene expression, considering that the expression of ARG1 in normal bone, inflammatory tissue, and 6 month post-surgery tissue at the healing site and the THP-1 derived macrophage populations was either very low or too low for detection. Detection of the expression of ARG1 at the gene and protein levels failed. Whether ARG1 could serve as a marker for human macrophages to classify inflammatory (M1) and anti-inflammatory (M2) macrophages needs further exploration. It should be provided further data to illustrate the mechanism of ARG1 expression by the human M2 macrophages during material-mediated bone regeneration and the mechanism of representation of the repair effect via the expression of ARG1.

## 3 Methods

### 3.1 Retrieval of autogenous bone tissue, inflammatory bone tissue and bone at healing sites

All surgical procedures and post- operative care were under the approval of the Ethics Committee (WCHSIRB-D-2017-033-R1), and were performed at the Implantology Department from 2017 to 2018. After tooth extraction, the inflammatory periodontal tissues within sockets were harvested (IP). During subsequent osteotomy, autogenous bone tissue was collected (NB). Guided bone regeneration was performed in the empty alveolar socket using deproteinized bovine bone mineral (DBBM, BioOss, Geistlich, Switzerland). Following a 6M healing period, a trephine (diameter of 3mm) was used to harvest the local bone block during implant site preparation (H). Those harvested tissues were gently washed in sterile saline (30 s) before placing in a cryopreservation tube (1.5ml) and stored in liquid nitrogen for subsequent testing, including RNA-seq, quantitative real-time polymerase chain reaction (qPCR) analysis, and flow cytometry analysis, while stored in 4% paraformaldehyde at 4°C for immunofluorescent staining. About the guided bone regeneration surgery, it began with minimally invasive tooth extraction. Then deproteinized bovine bone mineral particles (DBBM) were implanted to guide bone regeneration. At 6M post-surgery, bone tissue was harvested for RNA-seq, qPCR and flow cytometry analysis.

### 3.2 RNA-seq analysis

The bulk-seq analysis was carried out by Novogene Corporation (China). RNA was extracted from tissues or cells using standard methods to make sure samples were strictly controlled for quality. The standard procedure mainly included the following three aspects: analysis of sample RNA integrity, DNA contamination and detection of RNA purity (OD260/280 and OD260/230). In terms of library construction and quality control, mRNA can be obtained in two main ways: firstly, most eukaryotes’ mRNA has poly A-tailed structural, and poly A-tailed mRNA can be enriched by Oligo (dT) magnetic beads. The other is the removal of ribosomal RNA from the total RNA to obtain mRNA. Subsequently, the obtained mRNA was randomly interrupted by divalent cations in NEB Fragmentation Buffer, and the database was constructed according to the NEB general database construction method or chain specific database construction method. Upon completion of library construction, a Qubit2.0 Fluorometer was used for initial quantification, and the library was diluted to 1.5ng/ul. Then the insert size of the library was detected using Agilent 2100 bioanalyzer. After the insert size met the expectation, the effective concentration of the library was accurately quantified by qRT-PCR (the effective concentration of the library higher than 2nM) to ensure library quality. Finally, the libraries were qualified for sequencing, and Illumina sequencing was performed after pooling the different libraries according to the requirements of effective concentration and target data volume, of which the basic principle is sequencing by Synthesis. Gene count and Fragments Per Kilobase of transcript per Million mapped reads (fpkm) of the selected genes were showing in the bar chart (Fig. 1b and 1c).

### 3.3 Reagents and materials

All cell culture reagents were purchased from Life Technologies (USA), unless otherwise noted. Antibodies: PE-conjugated anti-mouse iNOS (Cat. #NBP2-22119, Clone 4E5) was purchased from Novus Biologicals (USA). FITC-conjugated anti-mouse CD68 (Cat. #ab134351, Clone Y1/82A) and PerCP/Cy5.5®-conjugated anti- rat CD11b (Cat. #ab210299, Clone M1/70) were purchased from Abcam (UK). PE/Cyanine-conjugated anti-mouse ARG1 (Cat. # 369707, Clone 14D2C43) was purchased from BioLegend Inc. (USA). HyClone phosphate buffered saline (PBS, Cat. #SH30256.01) and RPMI medium modified (Cat. #SH30809.01) were purchased from GE Healthcare Life Sciences HyClone Laboratories (USA). Bovine serum albumin V (BSA V, Cat. #A8020-5g) was purchased from Solarbio (China). Trypsin-EDTA (0.25%), phenol red (Cat. #25200056) and Gibco fetal bovine serum (FBS, Cat. #10099141C) were purchased from Thermo Fisher Scientific (Waltham, USA). Falcon® 70 μm Cell Strainer (Cat. #352350) was purchased from Corning (USA). PrimeScript™ RT Reagent Kit with gDNA Eraser (Perfect Real Time, Cat. #RR047A) and TB Green® Premix Ex Taq ™ (Tli RNaseH Plus, Cat. # RR402A) were purchased from Takara Bio Inc. (Japan).

### 3.4 Cell Culture (THP-1 monocytes derived macrophages’ response to different stimuli)

In order to verify the expression of ARG1 in macrophages described in the discussed literature, we consulted the original THP-1 monocytes (Procell, China) culture process and made some necessary changes as the anti-inflammatory/pro-inflammatory environment was simulated by IL4/LPS stimulation, and a group of common bone graft materials, DBBM, was added. To obtain macrophage, RPMI-1640 containing 100ng/ml phorbol myristate acetate (PMA) and 10%FBS (Gibco, USA) was treated to induce human THP-1 monocytes differentiation at 37°C for 24 h. The entire process started with cell resuscitation, that is, the frozen cells were transferred to a 37°C water bath until no ice crystals remain, centrifuged at 1000rpm for 3 min, and then resuspended with RPMI-1640 (PM150110) medium containing 10% FBS (164210-500) and 0.05Mm β-mercaptoethanol (PB180633) along with 1% P/S (PB180120, Cat. #CM-0233, Procell) and incubated in the T25 cell culture flasks at 37°C and 5% CO2 for 48 h. After flow cytometry was performed to detect the ratio of macrophages, cells were cultured in 6-well plates for 7 days in different circumstances of RPMI which is alone or added respectively with 10% FBS and 10ng/ml IL4; 10% FBS and 10ng/ml LPS; or 10% FBS and 0.03g/well DBBM to detect polarization of macrophage.

### 3.5 Quantitative Real-Time Polymerase Chain Reaction (qPCR)

Total RNA of THP-1 monocytes derived macrophages cultured in different circumstance for 7 days was digested from 6-well plates using trypsin and then TRIzol™ Reagent (Cat. #15596026, ThermoFisher Scientific, USA) was added. The concentration and ratio of total RNA were detected by NanoPhotometer NP80 (Implen, USA) at wavelength of 260 nm and 280 nm. The cDNAs were synthesized using PrimeScript™ RT reagent Kit with gDNA Eraser (Perfect Real Time, Cat. #RR047A), then amplified by qPCR with the specific primers (Supplementary Table 1). PCR was performed on QuantStudio 3 Real-Time PCR Systems (ThermoFisher Scientific, USA). Each 20 μL of PCR mixture contained 10μl of TB Green Premix Ex Taq (Ti RNaseH Plus, 2X), 0.4μL of PCR Forward Primer (10μM), 0.4μl of PCR Reverse Primer (10μM), 0.4μl of ROX Reference Dye (50X), 2μl Template and 6.8μl of Sterile purified water. Samples were incubated at 1 cycle of 95°C for 30 s followed by 40 cycles of 95°C for 5 s and 60°C for 34 s, and ended up with a cycle composing of 95°C for 15 s, 60°C for 1 min and 95°C for 15 s. Results were analyzed using the comparative CT (2^-ΔΔCT) method to calculate gene expression fold changes normalized to the levels of Actin/Gapdh gene transcripts. The experiments were repeated for three times independently (n=3).

### 3.6 Flow cytometry analysis

The surface markers of macrophages and their phenotypes were examined by flow cytometry to ensure the turning of THP-1 monocytes to macrophages and to exam the polarization of THP-1 monocytes derived macrophages after culturing in different circumstances: RPMI with 10% FBS, RPMI with 10% FBS and 10ng/ml IL4, RPMI with 10% FBS and 10ng/ml LPS, RPMI with 10% FBS and 0.03g/well DBBM. Cells were isolated by trypsinization after cultured for 1 (THP-1 monocytes to macrophages) and 7 (THP-1 monocytes derived macrophages after culturing in different circumstances) days respectively, co-incubated with antibodies against iNOS (iNOS-PE, Clone 4E5, Novus Biologicals, USA), CD68 (CD68-FITC, Clone Y1/82A, Abcam, UK), CD11b (PerCP/Cy5.5®, Clone M1/70, Abcam, UK) and ARG1 (ARG1-PE/Cyanine, Clone 14D2C43, BioLegend, USA) at 1:400 dilution in the dark for 1 h at 4°C (100μl per antibody for each sample). All samples were centrifuged at 450RCF for 5 min at 4°C. Supernatants were removed by aspiration, 1ml 1×PBS solution containing 0.04% bovine serum albumin (BSA) was used to wash the cells for twice. Each sample was resuspended in 1ml of 4% paraformaldehyde, and the eventual FACS analysis was performed on NovoCyte Flow Cytometers (ACEA Biosciences®, USA) and FlowJo 10.5.0. The experiments were repeated for three times independently (n=3).

### 3.7 Immunofluorescent and Hoechst staining

After cultured for 7 days in different circumstances, cells were collected from 6-well plates and fixed in 4% paraformaldehyde solution for 24 h and washed 3 times in PBS. Then the sections were pretreated with 1% bovine serum albumin in PBS containing 0.1% Triton X 100 for 1 h, incubated in 1% Tween 20 for 20 min and washed again in PBS. The sections were subsequently analyzed for ARG1, according to the manufacturers’ instructions. Briefly, sections were incubated for 30 min in dark. The excess dye was rinsed off with PBS. Sections were incubated with antibody isotype to exclude false positive staining. At least three parallel sections from different implantation sites were observed with fluorescence microscope (ZEISS SteREO Discovery.V20, Germany). Five random sights were selected for immunofluorescence assay. Macrophages were observed at a magnification of 400 to exclude any false positive staining. Then semiquantitative analysis was conducted at a magnification of 40. The fluorescence intensity measurement was conducted according to five random sights with CaseViewer 2.1 and Image Pro Plus 7.0 (n=5). Meanwhile, the Hoechst staining were also performed for analyzation.

### 3.8 Statistical Analysis

Statistical significance for in vivo and in vitro data of qPCR or FACS were analyzed either by Student’s t-test, or one-way analysis of variance (ANOVA) at the 95% confidence level with Tukey’s post hoc test, which were performed in GraphPad Prism 8.0 (GraphPad Software, USA) and P < 0.05 was considered statistically significant, while P > 0.05 was considered having no statistical differences, which was marked with NS.

## 4 Author contribution

The work allocation has been readjusted as CHENYU CHU, CHEN HU and SHENGAN RUNG was mainly responsible for operating of subsequence experiments and revising whole manuscript. YUFEI WANG was mainly responsible for graphing the figures and clearing the thought and assistance of animal experiment and sample harvest. YILI QU and YI MAN was mainly responsible for management, design of experiment and surgical procedure. All authors agree with adjustment of order.

## 5 Acknowledgments

Funding information: National Natural Science Foundation of China, Grant/Award Number: 81870801 and 81671023.

---

**Supplementary figure 1.**
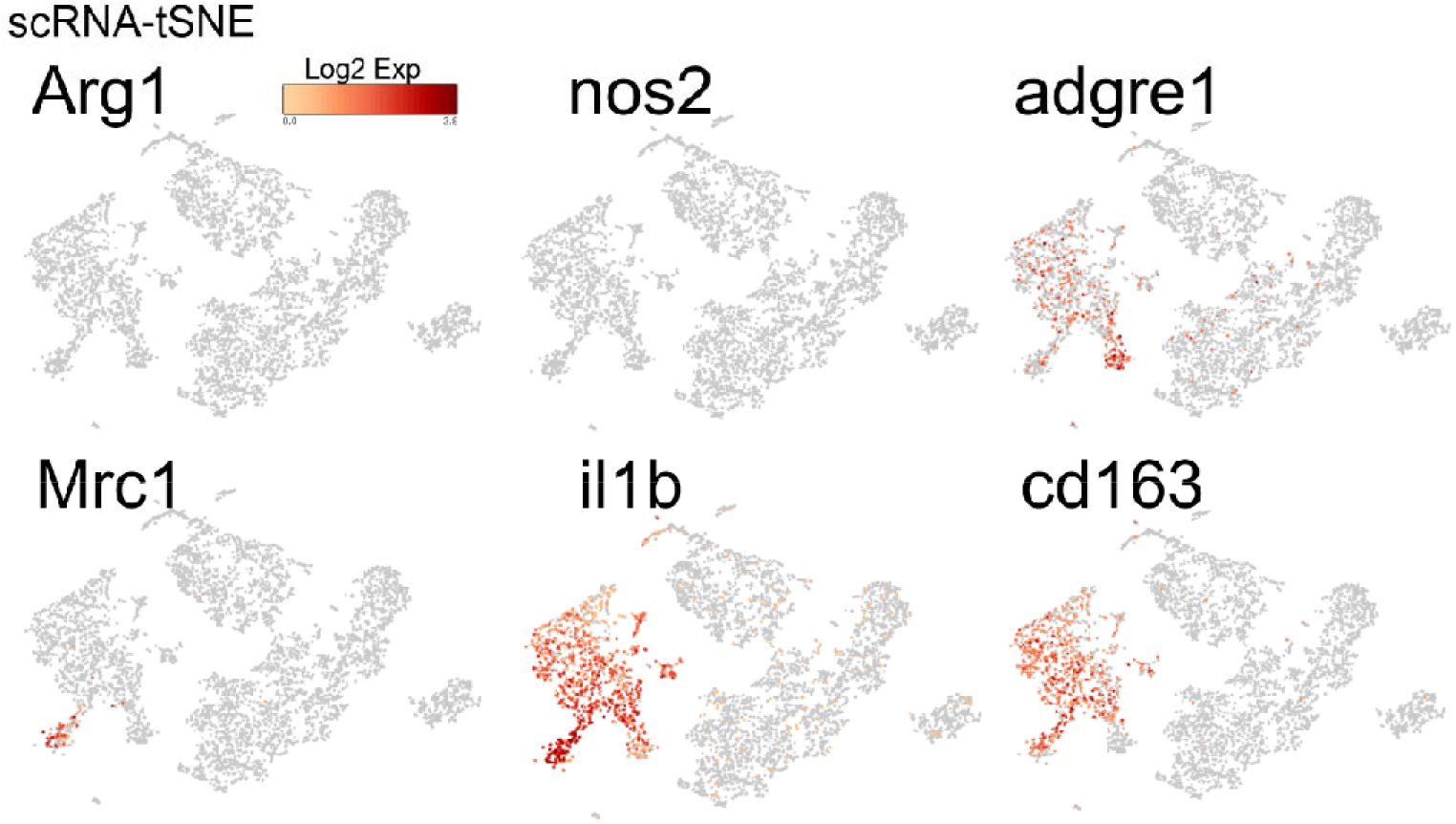
Performing single cell RNA sequencing on peripheral blood mononuclear cell (PBMC) of healthy human, ARG1 was not detected in macrophage-related genes.

**Supplementary figure 2.**
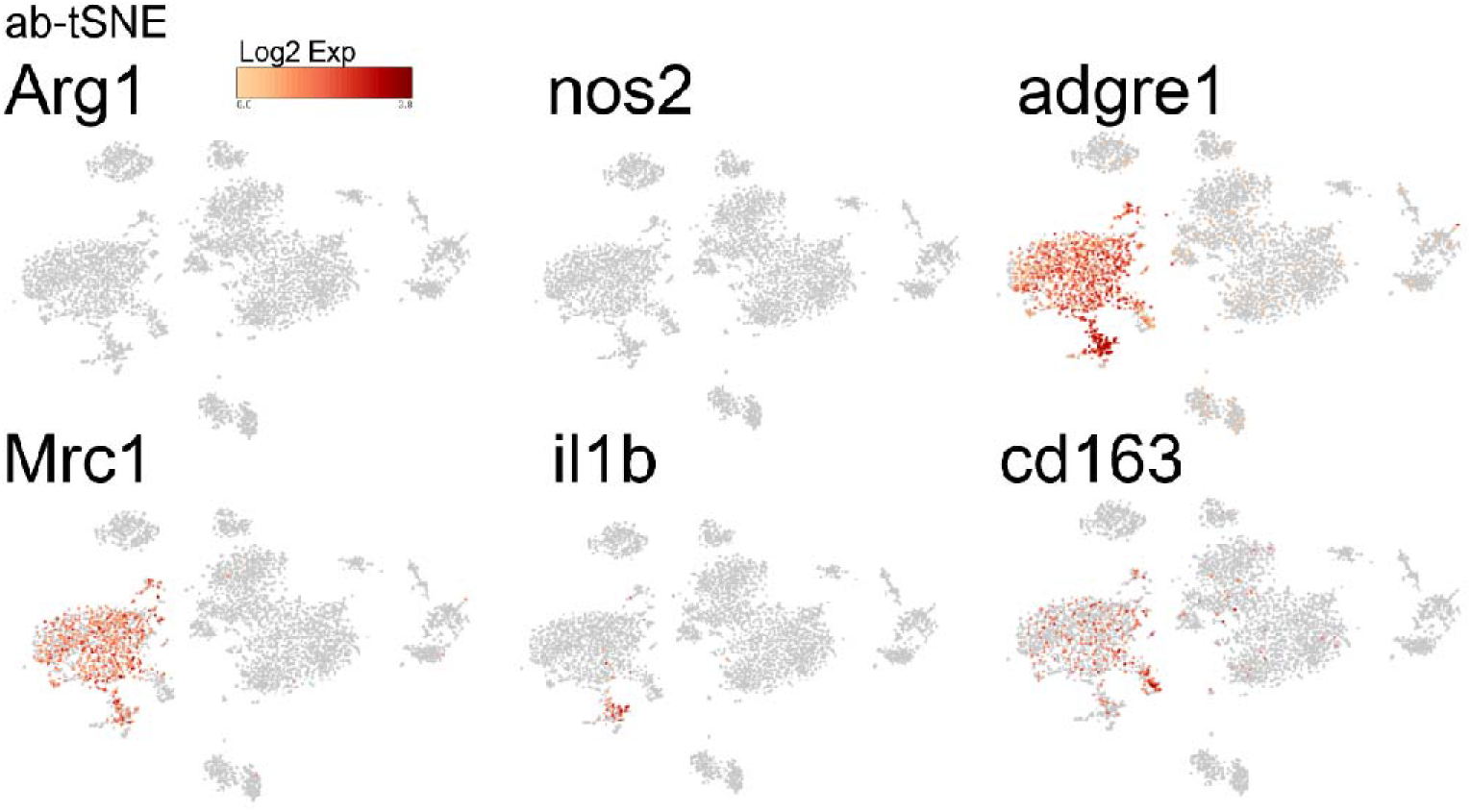
The antibody staining of markers of peripheral blood mononuclear cell (PBMC) showed that except ARG1 and NOS2, other macrophage-related genes were detected.

**Supplementary Table 1.**
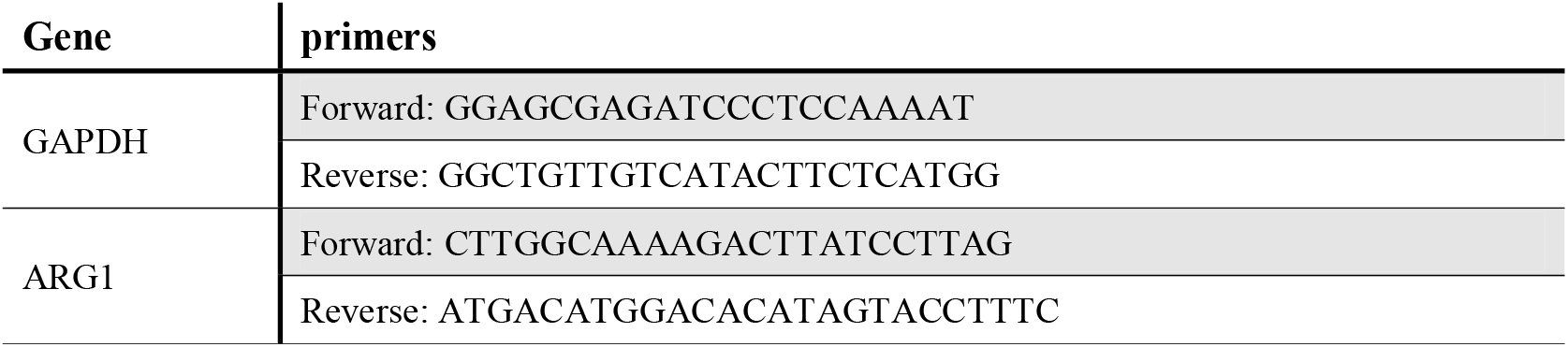
Forward and reverse primers corresponding to gene for qPCR analysis.

